# Brain-Computer Interfaces for Post-Stroke Motor Rehabilitation: A Meta-Analysis

**DOI:** 10.1101/224618

**Authors:** Maria A. Cervera, Surjo R. Soekadar, Junichi Ushiba, José del R. Millán, Meigen Liu, Niels Birbaumer, Gangadhar Garipelli

## Abstract

**Objective:** Brain-computer interfaces (BCIs) can provide sensory feedback of ongoing brain oscillations enabling stroke survivors to modulate their sensorimotor rhythms purposefully. A number of recent clinical studies indicate that repeated use of such BCIs might trigger neurological recovery and hence improvement in motor function. Here we provide a first meta-analysis evaluating the clinical effectiveness of BCI-based post-stroke motor rehabilitation.

**Methods:** Trials were identified using MEDLINE, CENTRAL, PEDro and by inspection of references in several review articles. We selected randomized controlled trials that used BCIs for post-stroke motor rehabilitation and provided motor impairment scores before and after the intervention. A random-effects inverse variance method was used to calculate the summary effect size.

**Results:** We initially identified 524 articles and, after removing duplicates, we screened titles and abstracts of 473 articles. We found 26 articles corresponding to BCI clinical trials, of these, there were nine studies that involved a total of 235 post-stroke survivors fulfilling the inclusion criterion (randomized controlled trials that examined motor performance as an outcome measure) for the meta-analysis. Motor improvements, mostly quantified by the upper limb Fugl-Meyer Assessment (FMA-UE), exceeded the minimal clinical important difference (MCID=5.25) in six BCI studies, while such improvement was reached only in three control groups. Overall, the BCI training was associated with a standardized mean difference (SMD) of 0.79 (95% CI: 0.37 to 1.20) in FMA-UE compared to control conditions, which is in the range of medium to large summary effect size. In addition, several studies indicated BCI-induced functional and structural neuroplasticity at a sub-clinical level.

**Interpretation:** We found a medium to large effect size of BCI therapy compared to controls. This suggests that BCI technology might be an effective intervention for post-stroke upper limb rehabilitation. However, more studies with larger sample size are required to increase the reliability of these results.

## INTRODUCTION

Stroke is the second leading cause of death worldwide, with 6.7 million cases registered in 2012.^1^ It is also one of the leading causes of disability with an estimated 50% of the survivors suffering from permanent motor or cognitive impairments.^2^ Upper limb disability is particularly critical as it is highly prevalent and vastly reduces the independence in activities of daily living (ADL).^3,4^ Currently, motor rehabilitation techniques for stroke patients with hemiplegia usually include physical therapy and constraint-induced movement therapy (CIMT),^5^ which require some residual movement of the affected limb. However, approximately 20-30% of all stroke survivors do not qualify for CIMT or other rehabilitation strategies. For those patients, mirror therapy,^6^ motor imagery,^7,8^ action observation therapy,^9^ electrical stimulation (e.g., non-invasive brain stimulation,^10–12^ or vagus nerve stimulation,^13^) and robot-aided sensorimotor stimulation^14^ have been investigated as possible alternatives over the last several years. Driven by advances in other technological areas such as virtual and augmented reality (VR/AR), robotics, invasive and non-invasive brain-computer interfaces (BCIs),^15^ as well as pharmacology,^16,17^ post-stroke motor rehabilitation is now a fast growing, emerging field.

A BCI translates electric, magnetic or metabolic brain activity into control signals of external devices that may replace, restore, enhance, supplement or improve the natural neural output, and thereby changes the ongoing interaction between the brain and its external or internal environment.^18^ A BCI can be invasive or non-invasive based on its brain activity measurement methodology. In invasive systems, electrodes are positioned on the surface of the brain (electrocorticography or ECoG) or implanted into the cortex (microelectrode arrays). In non-invasive systems, electrodes are placed on the scalp (electroencephalography or EEG, near-infrared spectroscopy or NIRS). In a typical EEG-based non-invasive BCI, user’s movement intention (motor imagery or execution) is decoded in real-time from the ongoing electrical activity of the brain by extracting relevant features (Figure 1). In a typical trial, the detection of movement intention would trigger a contingent sensory feedback to the user. This feedback can be delivered in an abstract form (e.g. a moving cursor on a computer screen) or as embodied feedback (e.g. visual representations of the participants body parts over a virtual avatar on a computer screen, in a VR head-mounted display or directly overlaid on the participant’s limbs; or somatosensory representations delivered through robotic, haptic or Neuromuscular Electrical Stimulation (NMES) systems) reproducing the intended movement, which was shown to enhance motor learning.^19–21^

**Figure 1.**
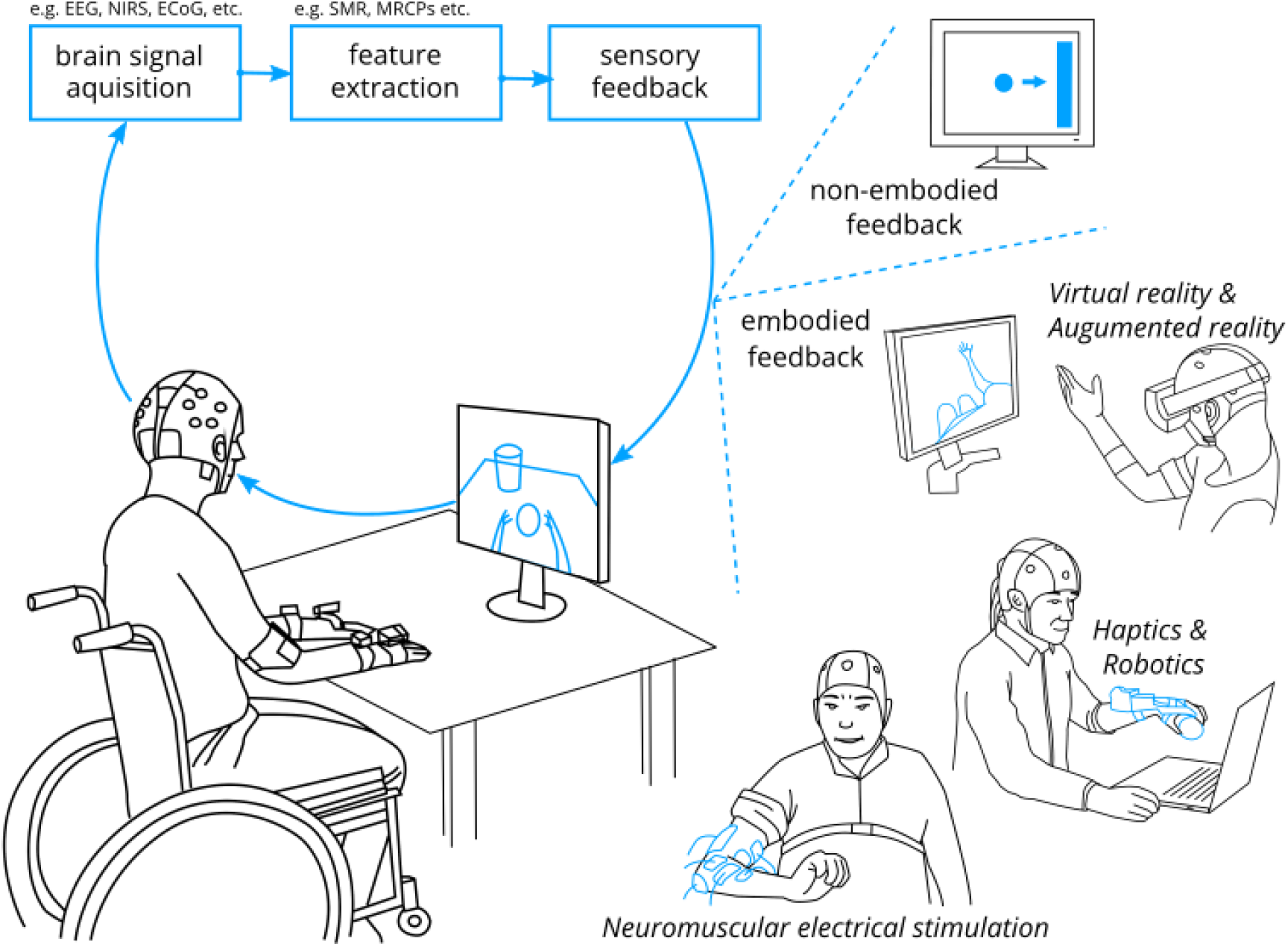
Illustration of typical brain-computer interface (BCI) systems used in post-stroke motor rehabilitation highlighting sensory feedback modalities. EEG = electroencephalography, NIRS = near-infrared spectroscopy, ECoG = electrocorticography, SMR = sensorimotor rhythm, MRCP = motor-related cortical potential.

BCIs are currently mainly explored in two clinical applications: (i) Assistive technologies that aim to restore lost functions, e.g. communication in locked-in syndrome (LIS) (e.g., as a result of amyotrophic lateral sclerosis)^22^ or movements in paralysis e.g., eating and drinking despite quadriplegia in an everyday life environment (using robotic actuators and/or functional electrical stimulation systems).^23^ (ii) Rehabilitation technologies also referred to as neurofeedback or rehabilitative BCIs,^24^ that aim to foster neuroplasticity through manipulation or self-regulation of neurophysiological activity facilitating motor recovery. In the current work, we focus on BCIs as rehabilitative technology in post-stroke motor rehabilitation.

Depending on movement complexity (unilateral vs. bi-lateral^25^) and the proximity of the muscle groups to the sagittal plane of the body (shoulder vs. hand^26^), movement-related neural activity was found to be not only present in the contralateral side but also on the ipsilateral side. Although the role of the unaffected hemisphere in post-stroke recovery is unclear, the ipsilesional primary motor cortex (M1) is thought to play a major role in motor recovery. Typical BCI-based motor rehabilitation protocols have predominantly aimed at cortical reorganization of the lesioned hemisphere.^27^ Specifically, most BCI-based motor rehabilitation systems have traditionally encompassed neural activity decoders of ipsilesional sensorimotor activity (sensorimotor rhythm, SMR, 9-15 Hz). Interestingly, a few recent studies suggest promoting contralesional hemispheric activity in moderate-to-severe chronic post-stroke patients, with an assumption that it may be harder to measure stable SMR from the ipsilesional sensorimotor areas in this group of patients^28^. Depending on the study protocol, BCI-mediated training may promote activity in ipsilesional or contralesional hemisphere.^29^

The power decrease of SMR during an attempt to move the paralyzed limb was shown to be associated with an increase in the excitability of the motor cortex,^30,31^ disinhibition of GABAergic inhibitory interneurons,^32^ increased excitability of the corticospinal tract^33^ as well as spinal motoneuron pools.^34^ An associated real-time feedback system (e.g. a robotic orthosis, NMES or a virtual reality avatar) that reproduces intended action (e.g., finger extension) allows patients to purposefully control sensorimotor oscillations.^24^ Similar to motor learning mechanisms, BCI-mediated motor training is thought to involve Hebbian neuroplasticity, error-based learning, and reward-based learning.^35^

In 2008, Buch et al. showed that severely paralyzed chronic stroke patients could learn to control their ipsilesional SMR.^21^ Since then, an international effort took place to investigate whether repeated BCI training can lead to motor recovery. Several studies reported neurological and behavioral improvements, such as increased event-related desynchronization (ERD) of SMR in the ipsilesional hemisphere,^36,37^ changes of motor-related functional connectivity assessed by functional MRI;^38^ increased control of volitional electromyographic (EMG) activity of the paralyzed muscles,^37,39^ and learned control of the reanimated hand and arm.^32, 34–37^ These results have encouraged the use of BCI in post-stroke motor rehabilitation, but the clinical efficacy is unknown so far. In this article, we aim to quantify the effectiveness of BCI training in post-stroke rehabilitation through a meta-analysis on existing randomized controlled trials (RCTs) reporting changes in motor function between the beginning and the end of the intervention. In doing so, we reviewed all available reports on RCTs using such technique and providing pre- and post-intervention motor impairment scores for both the experimental and control groups, which consisted of standard therapy, robotic therapy, electrical stimulation, motor imagery, or sham BCI feedback.

## METHODS

### Search Strategy and Eligibility Criteria

We searched for articles in MEDLINE^1^, CENTRAL^2^ and PEDro^3^ databases using brain-computer/machine interface, stroke, rehabilitation and trial as keywords. To identify all current trials, we also examined the references of over 20 key review articles (as of December 2016). We follow the PRISMA guidelines for reporting systematic literature review and meta-analysis (ref. supplementary materials).^43^ We included RCTs, where participants underwent BCI intervention for post-stroke motor rehabilitation. We excluded studies in which (non-sham) BCI was part of the therapy in both experimental and control groups, and studies that did not provide motor impairment assessment scores pre- and post-intervention. We considered studies published in peer-reviewed conference proceedings and journals to maximize the number of included trials.

Articles retrieved by the search were screened by reading the title and abstract. Potentially eligible studies were then analyzed in full length. Eligibility of the studies was assessed independently by two authors and discussed later to resolve any disagreement. Studies providing pre- and post-intervention motor outcomes were considered for the systematic review, whereas only studies providing motor impairment scores (such as Fugl-Meyer Assessment (FMA)^44^) were considered for the meta-analysis.

### Meta-Analysis Method

For each study, two authors independently extracted the following information and analyzed risk of bias: 1) participants’ characteristics (including sample size, age, time from stroke, type of stroke and motor impairment); 2) inclusion/exclusion criteria for the trial; 3) characteristics of the intervention; 4) outcome measures considered and 5) type of control group. We contacted the investigators whenever some key piece of information was missing in the published report.

The intervention effect for each study was calculated as the standardized difference in means (SMD) of mean change in selected outcome measure between the experimental and the control group, based on Hedge’s equation with a correction for small studies.^45^ Heterogeneity in the intervention effect is inevitable as the included trials had differences in the study design. Hence, we performed a DerSimonian and Laird’s random-effect analysis^46^ to estimate the mean intervention effect and its 95% confidence interval (CI). We further computed the 95% prediction interval (PI) of the effect estimate dispersion across studies, the interval where the intervention effect of a new study will fall with 95% probability. Heterogeneity between studies was calculated using Higgins’ I^2^ statistic (0%: homogeneity; 50%: moderate heterogeneity; 100%: heterogeneity),^47,48^ indicating the percentage of variance that can be due to actual inter-study heterogeneity. The possible causes for the heterogeneity are explored using two subgroup analyses: (i) control group selection and (ii) participant’s post-stroke recovery phase.

We also assessed the possibility of publication bias by plotting the SMD against its precision, measured as the standard error (SE) of SMD. We then conducted Egger’s linear regression method to detect funnel plot asymmetry^49^ and determine whether studies with negative results are missing in the literature. All the analyses presented in this report were performed in using the *mais* software package of *StataIC 14*.^44, 45^

## RESULTS

### Search Results

524 articles were initially identified, and after duplicate removal, the titles and abstracts of 473 publications were screened. Out of these, 26 articles were designated as BCI post-stroke motor rehabilitation trials (Figure 2). Out of these 26, 12 articles were discarded based on the following exclusion criteria: (a) redundant report (n=6; that correspond to a clinical trial already reported in an included article), (b) no valid BCI control (n=5; e.g., BCI was also used as the control intervention), or (c) not provided motor score outcome (n=1). The remaining 14 articles were kept for the qualitative synthesis. Interestingly, most of them reported FMA scores before and after the intervention (9 upper limb and one lower limb trials) and all were non-invasive trials.

**Figure 2.**
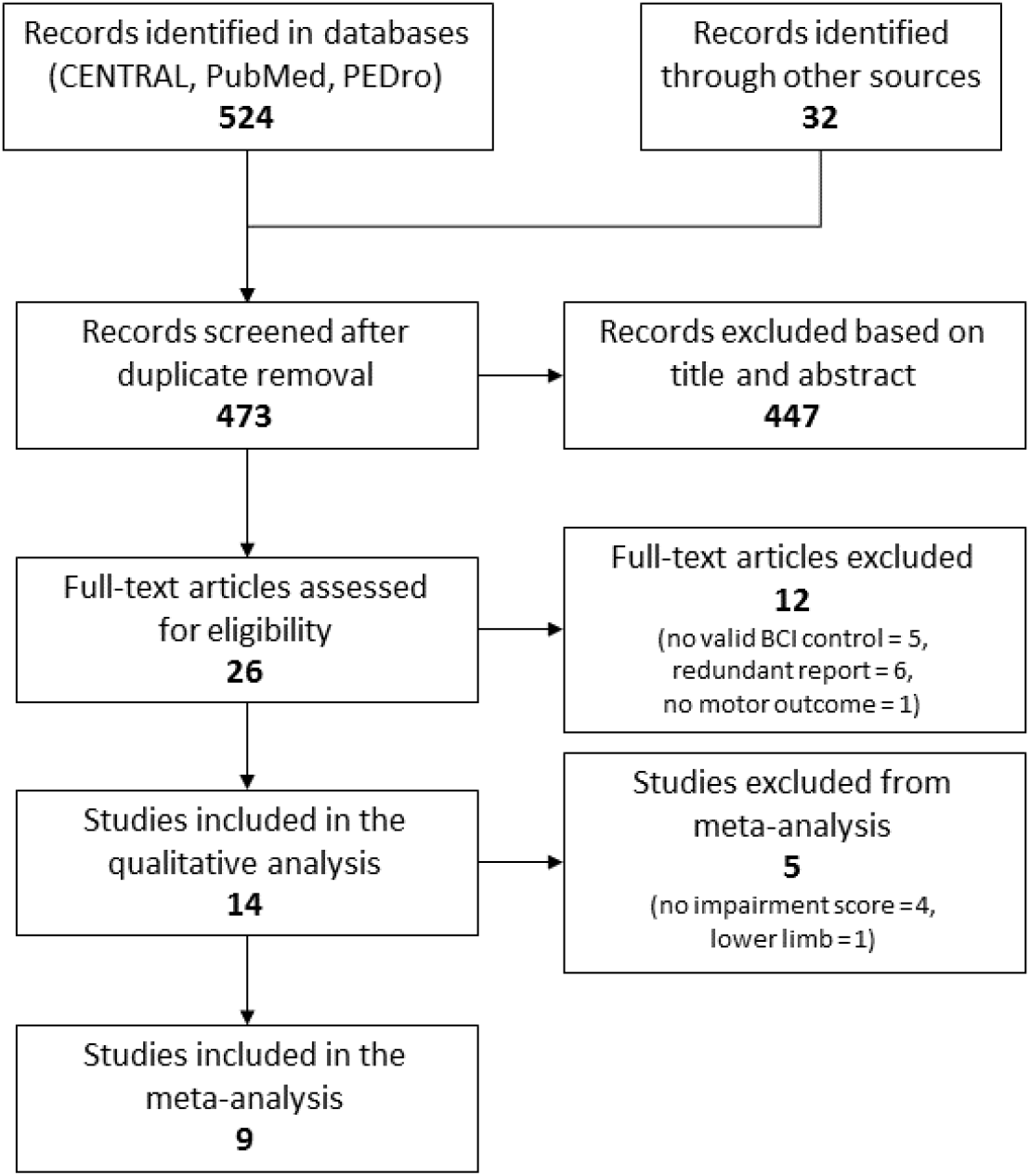
Flow diagram of study selection.

Among the 14 remaining trials, we found two upper limb studies that satisfied all inclusion criteria but did not measure FMA scores. The first, Rayegani et al. (N=20)^52^ reported significant improvement in Jebsen Hand Function Test^53^ score in six out of seven test items in the neurofeedback group and only three in the occupational therapy group. The second, Jang et al.^54^ compared a BCI system coupled to NMES with NMES alone for the treatment of shoulder subluxation in stroke patients (N=20). They reported a significant improvement in pain scores and Manual Function Test (MFT),^55^ but the difference between the groups was significant in only two items of the MFT.

We also found three studies that targeted lower limb: (i) Mrachacz-Kersting et al.^56^ reported improvements FMA for lower extremity (FMA-LE) score (mean difference of 0.8 ± 0.46) in chronic stroke patients (N=22), where they estimated reaction time from offline EEG, which was then used for delivering peroneal nerve stimulation during interventional trials (i.e. online trials), but not in the sham feedback group. Note that this study was not based on instantaneous decoding of movement intention for providing feedback, but was based on estimated reaction time from offline measurements. (ii) Chung et al. (N=10) reported significant differences in Timed Up and Go test, cadence and step length in the experimental group (BCI coupled NMES triggered ankle dorsiflexion) compared to a control group (NMES alone)^57^. Finally, Lee et al. (N=20) reported significant improvements in velocity and gait cadence in neurofeedback therapy compared to pseudo-neurofeedback control.^58^

We simplified the current study by restricting the meta-analysis to upper limb trials reporting FMA (total of 9 trials; excluding the above 2 upper limb trials and 3 lower limb trials from 14 studies), due to a limited number of available trials for other motor assessments and lower limb interventions.

### Characteristics of the Studies Included in the Meta-analysis

Among these nine studies (combined N=235, where sample size varied from 14 to 47; Table 1), one reported preliminary results of a clinical trial,^59^ and one reported results in a conference paper.^60^ Patients with first-ever ischemic or hemorrhagic stroke (cortical and sub-cortical) confirmed by a computer tomography or MRI scan and hemiplegia or hemiparesis caused by the stroke were included in these trials. Subjects were excluded if they had medical instability, cognitive or visual impairment, and high muscle spasticity. The mean age of the participants ranged from 49.3±12.5 to 67.1±5.51 years. Six studies targeted chronic patients,^39,41,59–62^ whereas the remaining three studies targeted,^42,63,64^ patients in the sub-acute phase, with a mean time from stroke approximately ranging from 2 to 4.5 months. In eight out of the nine studies, the BCI relied on the detection of ERD of SMR related to motor imagery. One study used near-infrared spectroscopy (NIRS) to measure task-related changes in levels of oxygenated and deoxygenated hemoglobin from the sensory-motor cortices.^63^ The motor intent detection signals were then used to trigger a sensory feedback provided by external devices (orthosis, robot, NMES system or visual display). The duration of therapies ranged from two to eight weeks. The nature of the control group differed across studies: sham-feedback triggered orthosis movement at random instances in four studies,^39,59,60,63^ one study used conventional therapy,^62^ one study used robot-assisted training,^41^ one study used NMES,^64^ and finally, one study used motor imagery.^42^ In ^61^, Ang et al. reported results of two different control groups: robot only and conventional therapy only. Whenever available, we decided to use the results of control groups undergoing conventional therapy. No significant adverse effects due to the rehabilitation were reported, although in one of the studies a patient dropped out due to a mild seizure during the intervention^61^.

**Table 1.**
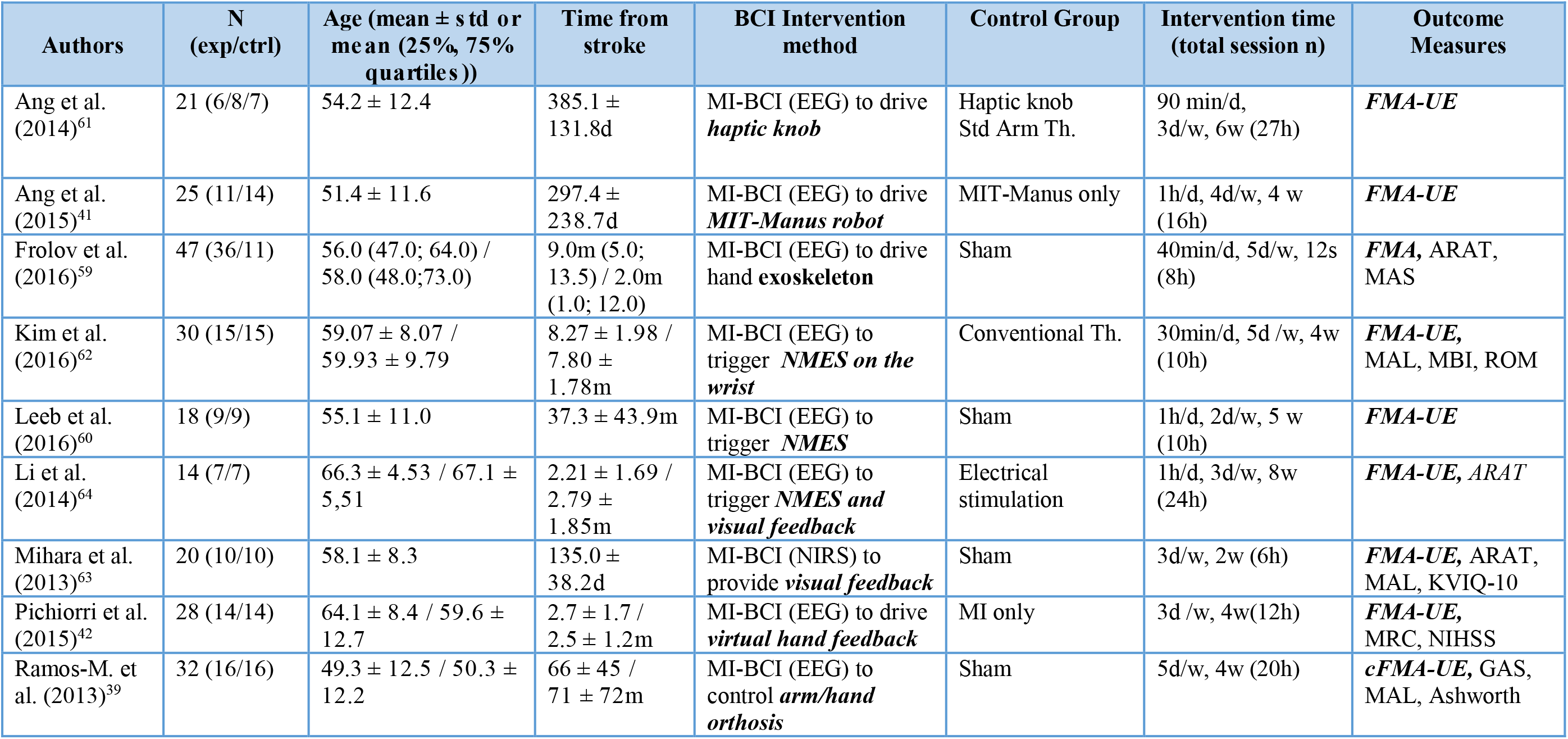
Characteristics of studies selected for the meta-analysis. Time from stroke and intervention characteristics are given in sessions (s) minutes (min), hours (h), days (d), weeks (w) and months (m). Provided outcome measures are: Fugl-Meyer Assessment for the Upper-Extremity (FMA-UE), Motor Activity Log (MAL), Modified Barthel Index (MBI), Range of Motion (ROM), Action Research Arm Test (ARAT), Kinesthetic and Visual Imagery Questionnaire-10 (KVIQ-10), Medical Research Council scale for muscle strength (MRC), National Institute Health Stroke Scale (NIHSS), Goal Attainment Score (GAS), Modified Ashworth Scale (MAS). Patient’s statistics (mean age and time from stroke) are provided either independently for the experimental and control groups or for the whole participant population depending on the data that was provided in each study.

| Authors | N (exp/ctrl) | Age (mean ± std or mean (25%, 75% quartiles)) | Time from stroke | BCI Intervention method | Control Group | Intervention time (total session n) | Outcome Measures |
| --- | --- | --- | --- | --- | --- | --- | --- |
| Ang et al. (2014)<sup>61</sup> | 21 (6/8/7) | 54.2 ± 12.4 | 385.1 ± 131.8d | MI-BCI (EEG) to drive *haptic knob* | Haptic knob Std Arm Th. | 90 min/d, 3d/w, 6w (27h) | *FMA-UE* |
| Ang et al. (2015)<sup>41</sup> | 25 (11/14) | 51.4 ± 11.6 | 297.4 ± 238.7d | MI-BCI (EEG) to drive *MIT-Manus robot* | MIT-Manus only | 1h/d, 4d/w, 4 w (16h) | *FMA-UE* |
| Frolov et al. (2016)<sup>59</sup> | 47 (36/11) | 56.0 (47.0; 64.0) / 58.0 (48.0;73.0) | 9.0m (5.0; 13.5) / 2.0m (1.0; 12.0) | MI-BCI (EEG) to drive hand *exoskeleton* | Sham | 40min/d, 5d/w, 12s (8h) | *FMA*, ARAT, MAS |
| Kim et al. (2016)<sup>62</sup> | 30 (15/15) | 59.07 ± 8.07 / 59.93 ± 9.79 | 8.27 ± 1.98 / 7.80 ± 1.78m | MI-BCI (EEG) to trigger *NMES on the wrist* | Conventional Th. | 30min/d, 5d /w, 4w (10h) | *FMA-UE*, MAL, MBI, ROM |
| Leeb et al. (2016)<sup>60</sup> | 18 (9/9) | 55.1 ± 11.0 | 37.3 ± 43.9m | MI-BCI (EEG) to trigger *NMES* | Sham | 1h/d, 2d/w, 5 w (10h) | *FMA-UE* |
| Li et al. (2014)<sup>64</sup> | 14 (7/7) | 66.3 ± 4.53 / 67.1 ± 5.51 | 2.21 ± 1.69 / 2.79 ± 1.85m | MI-BCI (EEG) to trigger *NMES and visual feedback* | Electrical stimulation | 1h/d, 3d/w, 8w (24h) | *FMA-UE*, ARAT |
| Mihara et al. (2013)<sup>63</sup> | 20 (10/10) | 58.1 ± 8.3 | 135.0 ± 38.2d | MI-BCI (NIRS) to provide *visual feedback* | Sham | 3d/w, 2w (6h) | *FMA-UE*, ARAT, MAL, KVIQ-10 |
| Pichiorri et al. (2015)<sup>42</sup> | 28 (14/14) | 64.1 ± 8.4 / 59.6 ± 12.7 | 2.7 ± 1.7 / 2.5 ± 1.2m | MI-BCI (EEG) to drive *virtual hand feedback* | MI only | 3d /w, 4w(12h) | *FMA-UE*, MRC, NIHSS |
| Ramos-M. et al. (2013)<sup>39</sup> | 32 (16/16) | 49.3 ± 12.5 / 50.3 ± 12.2 | 66 ± 45 / 71 ± 72m | MI-BCI (EEG) to control *arm/hand orthosis* | Sham | 5d/w, 4w (20h) | *cFMA-UE*, GAS, MAL, Ashworth |

Two authors independently assessed the risk of bias studies selected for the meta-analysis; disagreements were resolved by discussion. Six different factors proposed by the Cochrane Organization^4^ were analyzed in each study.(i) For each of these elements, authors assessed the risk as low (“+”), high (“-”) or unclear (“?”) following the Cochrane guidelines. (ii) Whenever information could not be found in the published reports, we contacted the authors for more details. A summary of the risk of bias under the six factors is illustrated in.

### Meta-Analysis of Upper Limb Intervention Trials

The mean and standard deviations of the FMA for upper extremity (FMA-UE) changes for the experimental and control groups in each study are presented in Table 3. The number of groups that showed improvements above minimal clinical important difference (MCID=5.25 ^65^) was six and three for BCI groups and controls respectively. The results of the main meta-analysis comparison are presented in a forest plot (Figure 3). The SMD favors BCI therapy versus control in eight out of nine studies. The most effective therapy was reported by Kim et al., where an SMD of 1.86 was found between BCI and control conventional therapy groups.(36) In five studies, the lower bound for the 95% CI lies above the no-effect (SMD=0) vertical line. The only result not favoring BCI was presented in Ang et al. with an SMD of −0.26.^41^ The combined intervention effect found is with an SMD of 0.79 (95% CI: 0.37 to 1.20). The weights of the studies, which are a function of the SE of the intervention effect, range from 8.45% to 14.00%; the contributions of each study to the result are comparable. Finally, we observed an *I*^2^ coefficient of 51.1%, reflecting considerable heterogeneity in the intervention effect. The 95% PI ranged from -0.39 to 1.97, showing that most new studies are likely to fall on the positive side, and a few are expected to report negative results.

**Figure 3.**
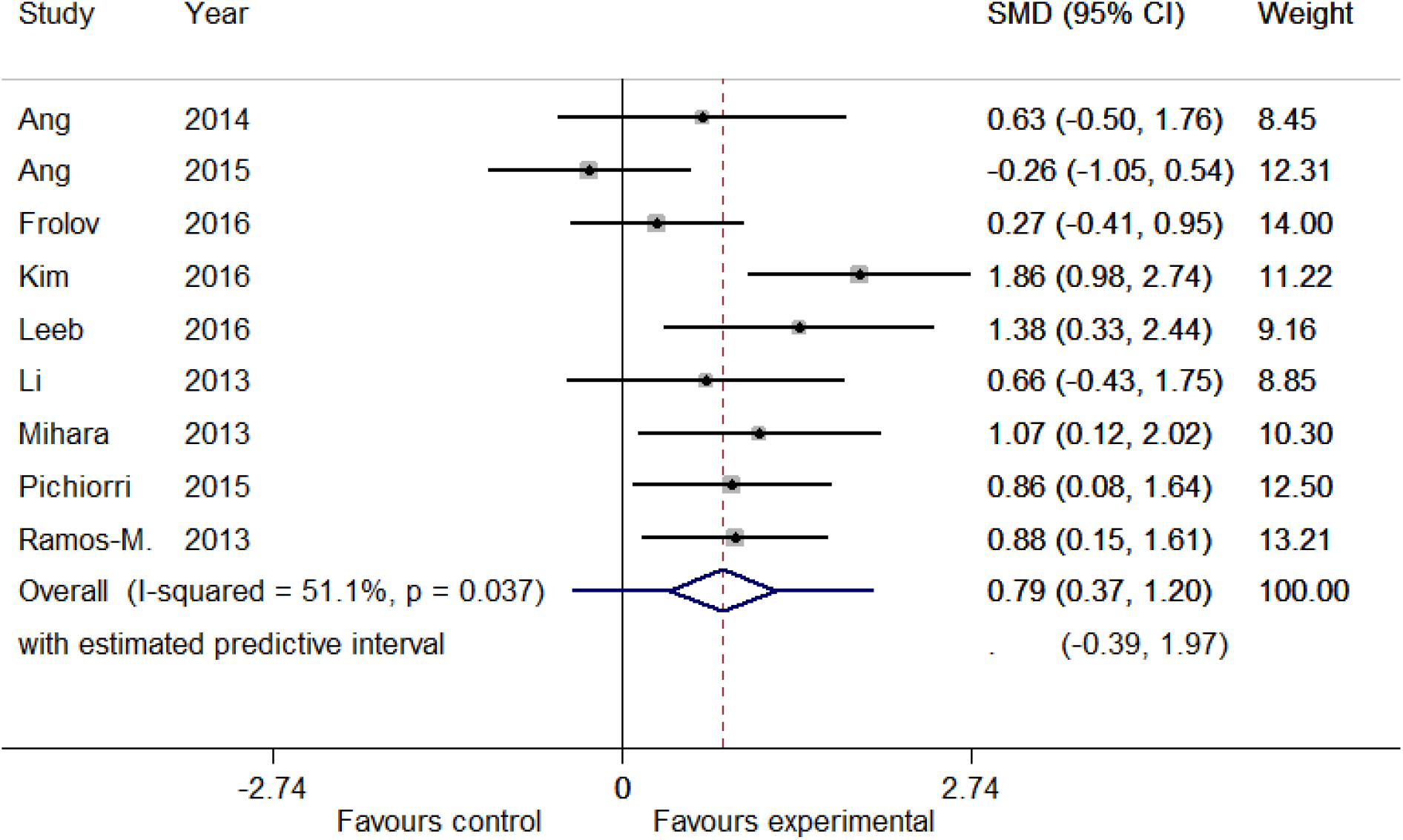
Intervention effect measured as changes in upper-extremity Fugl-Meyer Assessment (FMA-UE) scores between pre and post intervention (standardized mean difference (SMD), Random-Effects). The mean effect is represented as a diamond in the forest plot, whose width corresponds to the 95% CI, whereas the PI is shown as a bar superposed to the diamond. Box sizes reflect the contribution of the study towards the total intervention effect.

**Table 2.** Risk of bias of upper limb studies included in the meta-analysis (“+” = low risk; “-” = high risk; “?” = unclear risk).

| Study | Random Sequence Generation | Allocation Concealment | Blinding of participants and personnel | Blinding of outcome assessment | Incomplete outcome data | Selective reporting |
| --- | --- | --- | --- | --- | --- | --- |
| Ang et al. (2014)<sup>50</sup> | + | + | ? | + | + | + |
| Ang et al. (2015)<sup>36</sup> | + | + | ? | + | + | + |
| Frolov et al. (2016)<sup>48</sup> | ? | ? | + | + | + | + |
| Kim et al. (2016)<sup>51</sup> | + | + | + | + | + | + |
| Leeb et al. (2016)<sup>49</sup> | + | + | ? | ? | + | ? |
| Li et al. (2013)<sup>53</sup> | + | + | ? | ? | + | + |
| Mihara et al. (2013)<sup>52</sup> | + | + | + | + | + | + |
| Pichiorri et al. (2015)<sup>37</sup> | ? | + | + | + | + | + |
| Ramos-M et al. (2013)<sup>34</sup> | + | + | + | + | + | + |

**Table 3.** Mean FMA-UE changes (m) with standard deviations (sd) and number of subjects (n) in the BCI and control groups for the included upper limb trials.

| Study | Experimental |  |  | Control |  |  |
| --- | --- | --- | --- | --- | --- | --- |
| FMA-UE Change | m | sd | n | m | sd | n |
| Ang et al. (2014) <sup>61</sup> | 7.2 | 2.3 | 6 | 4.9 | 4.1 | 7 |
| Ang et al. (2015) <sup>41</sup> | 4.55 | 6.07 | 11 | 6.21 | 6.33 | 14 |
| Frolov et al. (2016) <sup>59</sup> | 5.25 | 4.50 | 36 | 4.09 | 2.91 | 11 |
| Kim et al. (2016) <sup>62</sup> | 7.87 | 2.42 | 15 | 2.93 | 2.74 | 15 |
| Leeb et al. (2016) <sup>60</sup> | 8.6 | 5.0 | 9 | 2.4 | 3.4 | 9 |
| Li et al. (2013) <sup>64</sup> | 12.7 | 11.3 | 7 | 6.7 | 4.1 | 7 |
| Mihara et al. (2013) <sup>63</sup> | 4.8 | 2.6 | 10 | 2.3 | 1.8 | 10 |
| Pichiorri et al. (2015) <sup>42</sup> | 13.6 | 8.9 | 14 | 6.5 | 7.0 | 14 |
| Ramos-Murguialday et al. (2013) <sup>39</sup> | 3.4 | 2.2 | 16 | 0.36 | 4.2 | 16 |

We found five sub-groups among all the control groups that may have impacted heterogeneity: (i) standard therapy, (ii) robot only, (iii) sham feedback, iv) NMES, and (v) motor imagery only (Figure 4). The smallest difference between experimental and control groups can be found for the robot only sub-group with an SMD of 0.63 (95% CI: - 0.50 to 1.76), whereas the major difference between the study arms is obtained for motor imagery sub-group where an SMD of 0.86 (95% CI: 0.08 to 1.64) was found. For the sham feedback sub-group, the intervention effect is slightly lower than for motor imagery sub-group, with an SMD of 0.80 (95% CI: 0.33 to 1.27). Overall, in all five sub-groups, the intervention was more effective in the BCI group compared to the control group. In the second subgroup analysis on the post-stroke recovery phase, we found higher intervention effect for the subacute sub-group with an SMD of 0.88 (95% CI: 0.35 to 1.41), compared to the chronic sub-group (SMD of 0.76 (95% CI: 0.15 to1.38)), but with a substantial overlap in CI (Figure 5).

**Figure 4.**
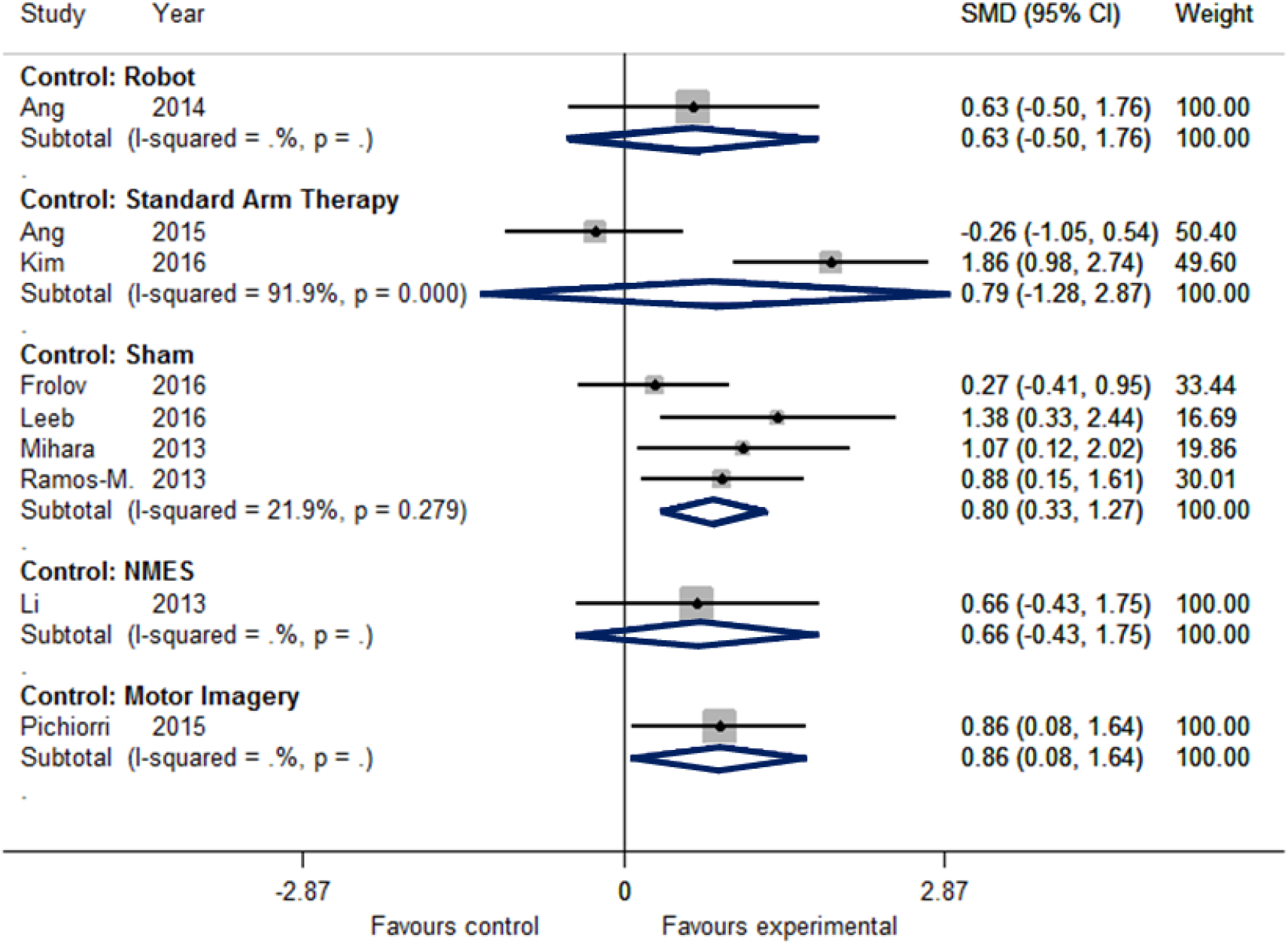
Subgroup Analysis 1: Standardized mean difference (SMD) of upper-extremity Fugl-Meyer Assessment (FMA-UE) scores in the studies under the random-effect assumption for the different interventions in the control group.

**Figure 5.**
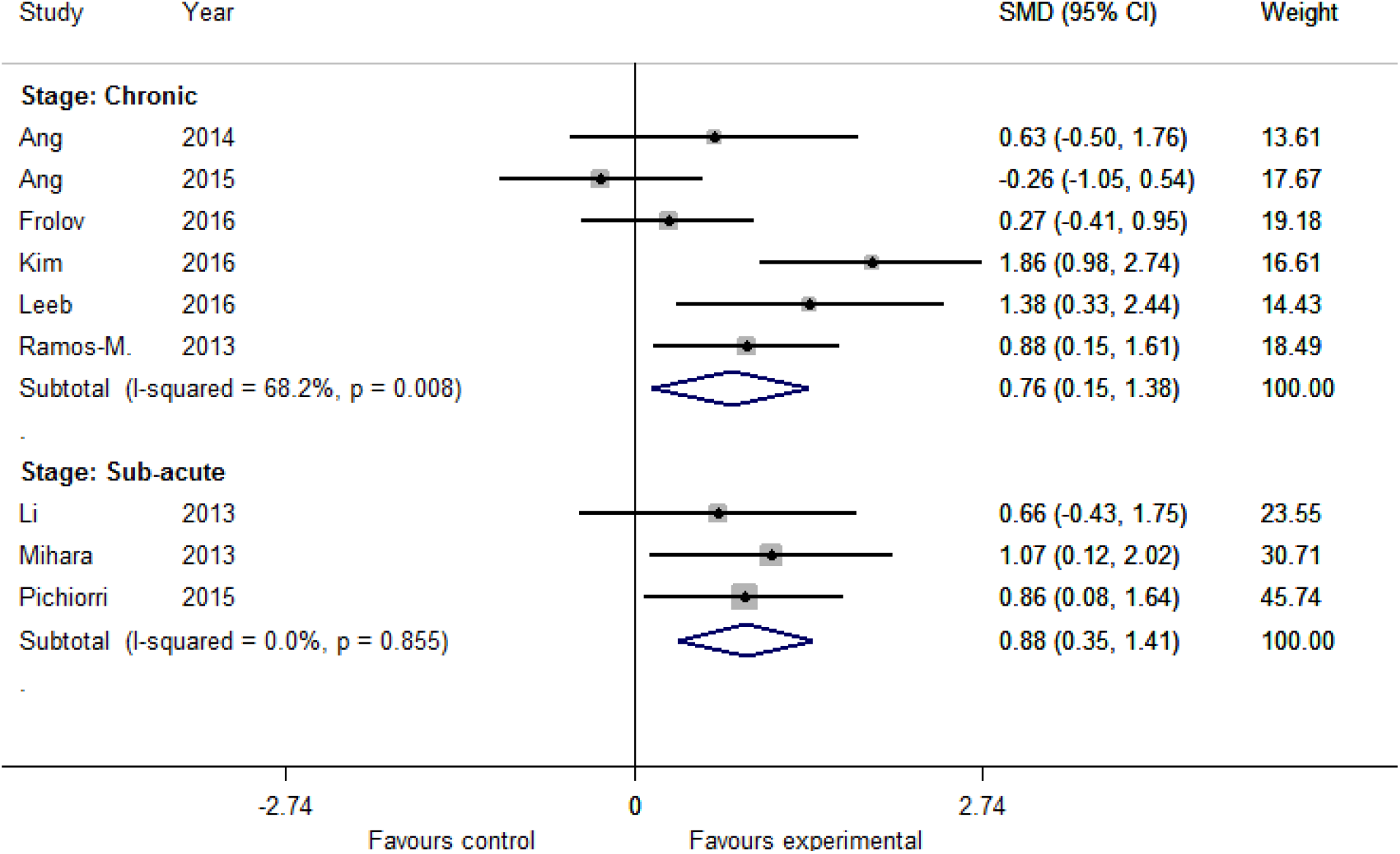
Subgroup Analysis 2: Standardized mean difference of upper-extremity Fugl-Meyer Assessment (FMA-UE) scores in the studies under the random-effect assumption. Studies are grouped into chronic and sub-acute phase.

We found no evidence of publication bias (Egger's test ^49^, p=0.353) by exploring the asymmetry of distribution of study findings and the summary effect size using a funnel plot (Figure 6). Studies reside at the bottom part of the plot, suggesting small sample sizes. Furthermore, two studies lie outside the region delimitated by the pseudo 95% confidence intervals (dotted lines), reflecting high heterogeneity between studies.

**Figure 6.**
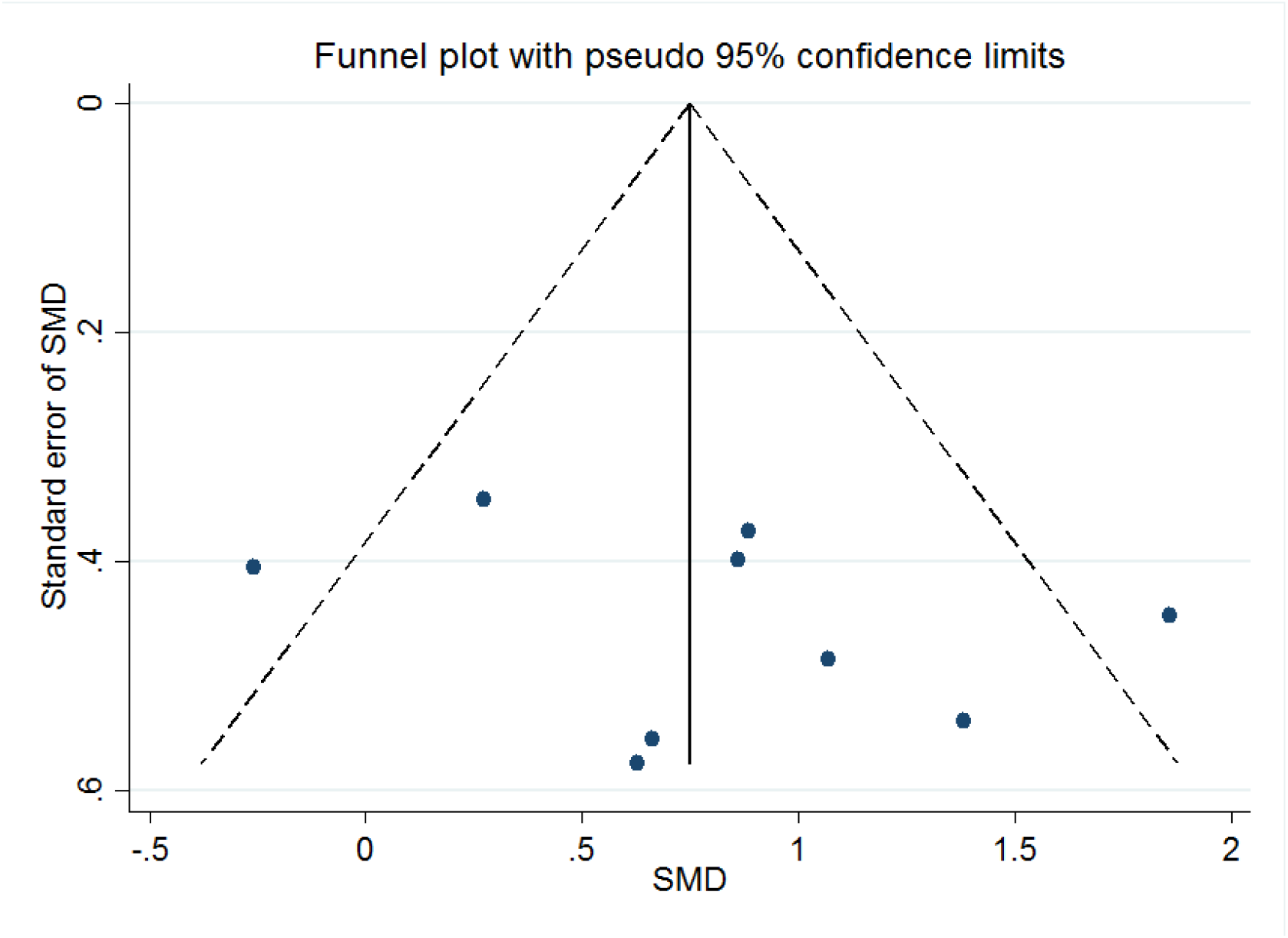
Funnel plot showing the precision (standard error, SE of standardized mean difference, SMD) against the effect size (SMD). The continuous vertical line shows the position of the overall combined effect, whereas dotted lines show pseudo 95% confidence limits.

## DISCUSSION

A number of clinical studies reported that repeated use of BCI systems after stroke could trigger neurological recovery, but the clinical effectiveness and effect size of repeated BCI-based neurorehabilitation training was unknown so far. We conducted a meta-analysis on available BCI intervention-based clinical trials as of December 2016. The analysis is limited to motor scales, as measurements of ADL and cost-effectiveness were unavailable. Due to a limited number of trials that reported non-FMA motor outcome measures and limited lower limb trials, we restricted the current meta-analysis to upper limb trials that reported FMA as a post-stroke motor outcome. A more comprehensive meta-analysis (e.g., examining the summary effect of various outcome measures such as motor/non-motor/ADL/stroke-severity and composite measures; as well as subgroup analyses such as early vs. late, upper limb vs. lower limb training and therapy dose) could be conducted in the future as the results of more randomized clinical trials become available.

### Interpretation of BCI intervention summary effect

Evaluating all available data on RCTs that applied BCI training to restore motor function after stroke, we found an SMD of 0.79 (95% CI: 0.37 to 1.20), meaning that the average FMA-UE score of the experimental group is separated by 79% of the pooled standard deviation from the control group. This evidence is in the medium (=0.35) to large (=0.8) range^66^ and comparable to widely applied therapy methods such as, CIMT (15 studies, N=355, SMD of 0.33 with 95% CI: 0.33 to 1.42, I^2^=78%)^67^ mirror therapy (11 studies, N=481, SMD of 0.61 with 95% CI: 0.22 to 1.00, I^2^=75%)^6^ and mental practice (5 studies, N=102, 0.62 of with 95% CI: 0.05 to 1.19).^68^ Furthermore, it clearly stands out in the context of other emerging technologies such as robotic interventions (31 studies, N=1078, SMD of 0.35 with 95% CI: 0.18 to 0.51, I^2^=36%)^6^, tDCS (7 studies, N=431, SMD of 0.11, with 95% CI of 0.33 to 1.42, I^2^=41%)^69^ and VR (10 studies, N=363, SMD of 0.27 with 95% CI: 0.05 to 0.49, I^2^=9%) interventions. Regarding actual FMA-UE scores, we found six studies with improvements that exceeded a MCID of 5.25 points in the BCI groups, whereas such improvements occurred in only three control groups. Of note, in six out of nine studies the differences in means of functional gains between the experimental and control groups remain below the MCID.^65^

The subgroup analysis of the type of control group revealed higher SMD for motor imagery than sham-feedback, robot, NMES and standard therapy. Similarly, we found a higher intervention effect for the sub-acute stroke group of studies with an SMD compared to the chronic stroke group. The subgroup analysis results are not conclusive though, due to the low number of studies that were included in each sub-group. There was no evidence of publication bias, but the included studies had low sample sizes.

### BCI-induced functional and structural neuroplasticity at a sub-clinical level

Since it was shown that repeated use of a neuroelectric or neuromagnetic BCI systems after stroke can lead to long-lasting effects on functional brain oscillatory activity (e.g., magnitude of event-related desynchronization^70^ or hemispheric blood-oxygen-level-dependent signals, BOLD^39^), follow-up studies indicated that such BCI paradigm may also lead to structural reorganization of the brain (as measured by diffusion tensor imaging^71–74^). Another study used real-time functional magnetic resonance imaging and showed that only two training sessions were sufficient to increase ipsilesional cortico-subcortical resting state connectivity in 3 out of 4 stroke survivors.^75^

This functional and structural re-organization may reflect improved motor planning and execution which, in some cases, may not have reached a measurable level using clinical assessments focusing on sensorimotor function (e.g., the FMA). Most published clinical trials using BCI systems for upper limb rehabilitation hint at such sub-clinical effects. Mihara et al. (2013)^63^, for instance, reported increased motor imagery-related BOLD activity in the pre-motor area in a group trained with a BCI compared to a group receiving placebo-BCI training. Similarly, Pichiorri et al. (2015)^42^ reported improved desynchronization in the mu and beta bands recorded over the ipsilesional primary motor cortex during motor imagery. Corbet et al.^60,76^ showed improved ipsilesional connectivity in the mu and beta bands after BCI-NMES training compared to a group undergoing placebo-BCI training. Other studies have reported shifts in hemispheric EEG activity^41^ and increase in ipsilesional movement-related cortical potentials (MRCP) as well as motor-evoked potentials (MEP)^56^. In addition, some of these neurophysiological measures correlated with behavioral improvements: (i) EEG-based Brain Symmetry index could predict the functional motor outcomes in Ang et al. (2015)^41^; (ii) changes in ipsilesional connectivity measures in mu-rhythms correlated with improvements in FMA-UL scores in both Corbet at al.^60,76^ and Pichiorri et al.^42^. In the future, BCIs could be further customized to facilitate structural and functional plasticity for re-organization of target brain regions and directed augmentation of sensorimotor maps to maximize their efficacy and viability in clinical applications.^77^

While the current results are encouraging, the field is yet to uncover the exact mechanisms of recovery underlying BCI training and the factors influencing BCI-aided rehabilitation success (e.g., type of lesion, the phase of recovery, dosage and intensity of BCI training). Major technological advances to maximize training effects, including the optimization of BCI system parameters, and to increase the practicality (e.g., shorter calibration) of these devices in a hospital or home environment are essential for the translation and broad adoption of BCI-based rehabilitation after stroke.

## CONCLUSIONS

Effects of BCI-based neurorehabilitation on upper-limb motor function show a medium to large effect size and can improve FMA-UE scores more than other conventional therapies. Besides motor outcomes, a number of studies also reported BCI-induced functional and structural neuroplasticity at a sub-clinical level, some of which also correlated with improved motor outcomes. More studies with larger sample size are required to increase the reliability of these results.

## Author Contribution

MC, NB, and GG are involved in data collection. MC and GG performed data analysis. MC, SRS, JU, JdRM, ML, NB and GG are involved in data interpretation, writing and editing the manuscript.

## Acknowledgements

With thank Mr. Ang, Ms. Mokienko, and Mr. Leeb for Ms. Chevalley for suggestions on the data collection and systematic review pipeline, Mr. Runnalls, Mr. Acon and Mr. Perez-Marcos for English proofreading, and finally Mr. Pardo for help with the illustration of Figure 1.

## Footnotes

1 https://www.ncbi.nlm.nih.gov/pubmed

2 http://onlinelibrary.wiley.com/advanced/search

3 https://search.pedro.org.au/search

4 http://www.cochrane.org/

